# Bioassay-guided isolation and characterization of anti-bacterial compound from *Sonneratia apetala* Buch. Ham leaf, from Maharashtra coast of India

**DOI:** 10.1101/2022.08.16.504112

**Authors:** Jaimini Dhane-Sarkar, Chiradeep Sarkar, B.L. Jadhav

## Abstract

Mangroves, collection of plants growing in coastal intertidal zones, are rich source of unique bioactive phytochemicals present in their bark, roots, leaves and fruits. They show anti-diabetic, anti-cancer, anti-hypertensive, anti-malarial, anti-bacterial, anti-inflammatory and anti-oxidant properties. In this study, a bioassay-guided isolation and characterization approach was taken to study the anti-bacterial compound from *Sonnerartia apetala* Buch. Ham leaf. Petroleum ether, acetone and water extracts were prepared and tested against eight human pathogenic bacteria. Water and acetone extracts have shown the inhibition of bacteria but the acetone extract was carried forward for further study. The compound was isolated using preparative column chromatography method. The isolated compound was studied for anti-bacterial activity using TLC-bioautography . The bioactive compound was isolated and tested against standard antibiotics and showed positive results against *E. coli* and *B*.*subtilis*. The isolated pure compound was studied for detection of elements and melting point. Further chacterization was done using UV-Vis spectroscopy and FT-IR spectroscopy indicating absorption in the UV spectrum and presence of aromatic and aliphatic groups. This study validates the importance of *Sonneratia apetala* leaves as an important resource in traditional medicine for anti-bacterial remedies aimed at human pathogenic diseases.

## Introdution

From last few decades, medical science is working on the challenges like malaria in general and specifically during pregnancy (Jaimini Sarkar, 2012; Gore-Langton et. al, 2022), Chronic Kidney Disease (Agarwal and Srivastava, 2009), Non-Alcoholic Fatty Liver Disease (Angulo and Lindor, 2002; Biju et. al, 2020; Bahirwani and Griffin, 2022), Polycystic Ovary Syndrome-PCOS (Rotterdam ESHRE/ASRM-Sponsored PCOS CWG, 2004; Sarkar and Maitra, 2013; Teede et al,2018), drug resistant TB (Udwadia et. al, 2011; Jaimini Sarkar, 2012), cancer incidence and deaths (Garcia et. al, 2007; Jaimini Sarkar, 2011) etc.

There are gender specific silent medical challenges like fibroids (Donnez & Dolmans, 2016; Jaimini Sarkar, 2013) and country specific neglected challenges like Human Rabies (Hemachudha et. al, 2002; Jaimini Sarkar, 2022; Gibson et. al, 2022) which need more attention.

Antimicrobial resistance to the known compounds (Ventola, 2015) and newly emerging diseases makes it essential to discover, design and engineer new promising classes of antibiotics (Gajdács, 2019).

Plants with ethnomedicinal significance when screened for phytoconstituents has shown compounds with diverse uses such as medicinal compunds (Abayomi, 1996; D. Jaimini & Jadhav, 2004; Mukharjee et. al, 2021), oils (Boukhatem & Setzer, 2020), food (Awuchi, 2019; D. Jaimini & Jadhav, 2004; D. Jaimini & Jadhav, 2005), fish meal supplements (Mallik et. al, 2019; Jadhav et. al, 2004; D. Jaimini & Jadhav, 2008).

Plants have been used in traditional medicinal practices since prehistoric times (Collins, 2000). They are valuable and indispensable sources of natural products with a great potential for producing new drugs for human beings (Nascimento et al., 2000; Morales et al., 2008; Littleton et al., 2005). Till date, number of phytochemicals with potential biological activity have been identified. In certain cases, the phytochemicals of some folklore medicinal plants and their pharmacological actions is still unexplored using scientific methods (Ahn, K., 2017). According to one estimate, only 20% of the plant flora has been screened for drugs (Parihar and Mathur, 2014). Consequently, their potential to be used as a new drug remains locked.

Many species of mangroves are being traditionally used to treat diseases such as rheumatism, smallpox, ulcers, hepatitis, leprosy, asthma, snake bites, toothache, and constipation (Prabhakaran & Kavitha, 2012). The mangroves have the ability to grow under stress factors such as high salt concentrations, tidal flooding, strong wind, solar radiations and heat (Spalding et al., 2010). This adaptability is based on morphological and physiological adaptations like stilt and air roots, salt excretion systems and high abundance of plant secondary metabolites (Glasenapp et al., 2019).

Medicinal potential of mangrove plants can be evaluated by the presence of their bioactive phytochemical constituents which shows activity like anti-inflammatory, anti-cancer, anti-viral, anti-malarial, anti-diabetic and anti-hypersensitive (Kathiresan & Bingham, 2001). However, since a single plant contains diverse phytochemicals, the effects of using a whole plant as a medicine are uncertain.

*Sonneratia apetala* is a fast growing (Chen et al., 2003), woody evergreen tree species of the mangrove (Mahmood, 2015). The *Sonneratia apetala* plant is commonly called as mangrove apple or kandel. The plant belongs to the family-Lythraceae, (Hossain et al., 2013). This tree species is native to Bangladesh, India, Sri-Lanka, Malaysia and Australia (Jayatissa et al., 2002; Liao et al., 2004). In local languages, it is known by various names like Kalinga in Telugu, Keruan in Oriya, Marama maram in Tamil, and Keora in Bengali (Shamina Nazrin et al., 2017).

*Sonneratia apetala* has immense medicinal and economic value. Various parts of this plant like bark, root, leaves and fruits have been used in folk medicine in the South Asian countries for treating diarrhoea, hepatitis, inflammation, wounds, ulcers (Hazra et al., 2021). The leaves of the plant are mainly used to treat hepatitis (Bandaranayake, 2002; Bandaranayake, 1998), for treating dysentery, sprains and bruises, in eye troubles and for open sores in children’s ears (Bandaranayake, 1995).

Few systematic studies have been carried out to evaluate the anti-microbial potential of *S. apetala* plant parts like leaves (Jaimini et al., 2011; Patra et al., 2015), bark (Md. Arifur Rahman et al., 2021) and fruits (Jana et al., 2015).

Scientific literature shows studies carried out for isolation and characterisation of phytochemicals from *S. apetala* leaf which reported presence of alkaloids, terpenoids, phenols (Sunita and Joshi, 2015) and phytosterols and Quinones (Ji. et al., 2005) from *S. apetala* leaf. But these attempts have not covered the anti-bacterial analysis of the isolated compounds.

There is need to isolate and characterize the specific phytochemicals with anti-bacterial potential from the *S. apeatala* plant leaf. A typical protocol to isolate a pure chemical agent from natural origin is bio-assay guided fractionation, meaning step-by-step separation of extracted components based on differences in their physico-chemical properties and assessing the biological activity, followed by next round of separation and assaying (Malviya and Malviya, 2017). With the objective of isolating compound(s) with anti-bacterial potential from *S. apetala* leaf, the current study has been designed using bio-assay guided isolation and characterization procedure.

## Materials and Methods

### Collection and Processing of Plant Material

Fresh, young and tender leaves of *S. apetala* Buch.-Ham (Lytheraceae) were collected from mangrove growing area of Ghodbunder Road, Thane, Maharashtra coast, India (Fig. I). Identity of the plant material was confirmed by an expert taxonomist of University of Mumbai, Mumbai. Leaves were dried in the shade at room temperature and dried leaves were ground to a fine powder in a Jankel and Kunkel model A10 mill. 10 gms of dried leaves powder was taken into cellulose thimble, single thickness supplied by Whattman’s International Ltd., Maidstone, England and placed carefully in the central tube of the extractor of the Soxhlet apparatus. The extraction was carried out using different polar and non-polar solvents (Petroleum Ether, Acetone and Water). The extract(s) was evaporated on water bath to a final volume of 20 ml to achieve the 50% (W/V) concentration. Later on, the extract was cooled and stored in airtight glass bottle at -4 °C in the refrigerator.

**Figure I.**
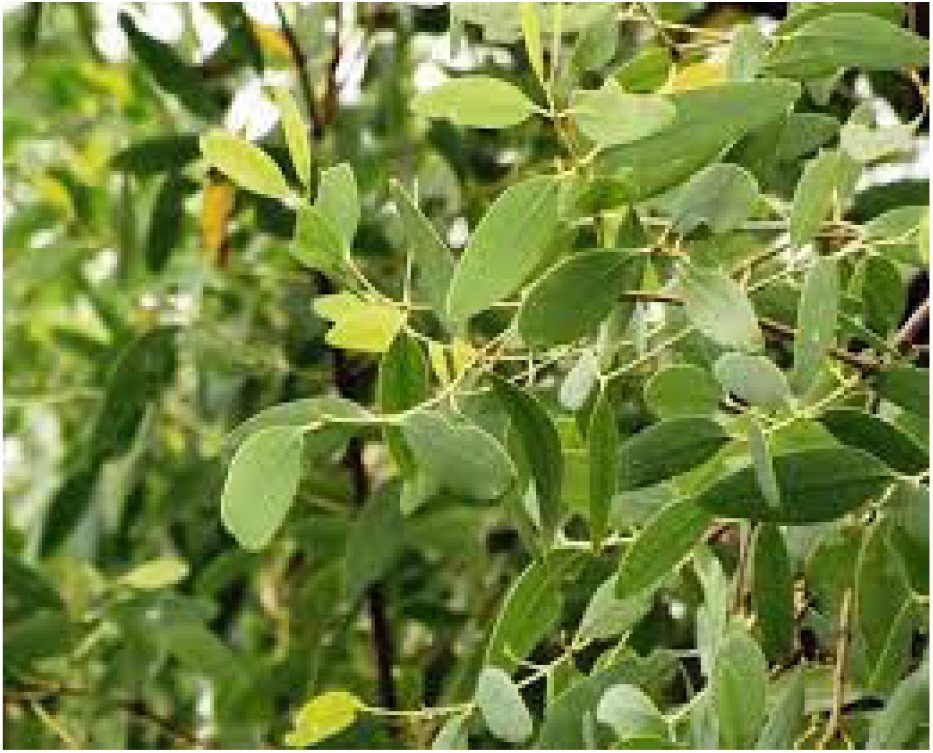
A typical *S. apetala* plant showing simple and broad leaves.

### Bioactivity evaluation of the extract

Extraction of *S. apetala* leaf was carried out using non-polar to polar solvents like petroleum ether, acetone and water. *In vitro* anti-bacterial activity of leaf extracts of *S. apetala* was carried out against eight human pathogenic bacteria. The bacterial species used included four Gram-positive (*Streptococcus pyogens, Staphylococcus aureus, Bacillus subtilis, Staphylococcus epidermidis*) and four Gram-negative bacteria (*Escherichia coli, Pseudomonas auriginosa, Salmonella typhi, Klebsiella aerogens*). These are clinical isolates from Haffkine’s Laboratory, (Mumbai) India. The anti-bacterial evaluation of the extract(s), MIC and bioactivity of pure compound was carried out against above mentioned bacteria by agar cup method (Spooner and Skyes, 1972).

Among petroleum ether, acetone and water, the petroleum ether extract did not show any anti-bacterial activity. Water and acetone extract has shown the inhibition of bacteria but the acetone extract was carried forward for the MIC. Compare to acetone(56 ^0^C), water (100 ^0^C) has higher boiling point. Consequently, it is not easy to prepare perfectly dried sample for MIC from water extract as compare to acetone extract. There is possibility that higher temperature will destroy some of the compounds of interest. At the same time, acetone has ability to dissolve both polar as well as non-polar substances, while other solvents can only dissolve one or the other (Ecolink, 2019). Considering all these reasons, acetone extract of *S. apetala* leaf was selected for the MIC study.

For MIC test, acetone extract was evaporated to dry residue and by using fresh solvent different concentrations (0.5, 1, 5, 10 mg/ml) of crude extracts were prepared.

Bioassay of pure compound was done at the concentration 250 μg/ml and compared with ten standard antibiotics (ampicillin, chloroamphenicol, cefazolin, tetracycline, minocyclin, vancomycin, sparfloxacin, erythromycin, tetracycline hydrochloride and benzylpenicillin sodium) against *E*.*coli* and *B. subtilis*.

### Development of solvent system

The solvent system(s) were developed using either individual solvent or mixture of solvents to obtain a range of polarity and appropriate phases for the better separation of the compound(s). The solvents used were petroleum ether, toluene, diethyl ether, chloroform, ethyl acetate, acetone, isopropanol, methanol and acetonitrile. In individual solvent system, diethyl ether, chloroform, ethyl acetate, and isopropanol has shown band(s) of separation. In combination of solvents, ten different combinations have shown band(s) of separation.

### Thin-layer chromatography-bioautography

TLC of potent crude extract(s) and fraction(s) isolated by preparative column chromatography was carried out by dipping method (Peifer, 1962). Single drop of sample was applied on TLC plate, about 2 cm from the edge with the help of capillary. The loaded TLC plate is allowed to develope in selected solvent system(s). Since many compounds separated by TLC are colorless, detection of separated bands on chromatoplates was carried out by exposing it to iodine vapor or placing in UV cabinet at 254 and 336 nm.

TLC-bioautography method introduced by Fisher and Lauther (1961) and Nicolaus et al., (1961) was carried out for the bioassay of the chromatoplates showing well separated band under UV. This method is also called as contact bioautography where anti-microbials diffuse from a TLC plate or paper to an inoculated agar plate. The chromatogram is placed face down onto the inoculated agar layer and left for some hours to enable diffusion. Then the chromatogram is removed and the agar layer is incubated. The inhibition zones are observed on the agar surface in the places where the spots of anti-microbials are stuck to the agar.

Contact bioautography of all the chromatoplates showing bands was carried out. Only one solvent system {Petroleum ether:methanol (7:3)} which showed one band was found to be active against *E. Coli* and *B. subtilis*. Hence, this particular solvent system was used for the preparative column chromatographic separation.

### Isolation of compound by preparative column chromatography

The glass column of length 75 cm and mouth with 2 cm diameter was rinsed with the help of petroleum ether and allowed to dry. Subsequently, column was packed with double the amount of alumina than the slurry. The residue of acetone fraction obtained by evaporating entire solvent was then added in alumina by continous and vigorous stirirng with the help of glass rod in a beaker, till the free flowing slurry was produced. Free flowing and thoroughly mixed slurry was placed on the top of the alumina. Once again, the cotton was placed on top of the slurry to avoid scattering of slurry/alumina in the solvent system. The solvent system {Petroleum ether:methanol (7:3)} was allowed to continuously pour on the column till the complete isolation of compound was achieved.

The purity of the compound was checked as a single TLC spot.

### Characterization of pure compound

Characterization of this organic compound was performed for solubility, detection of elements and melting point with standard laboratory tests.

### UV Spectroscopy

The dry, powdered compound was dissolved in methanol to make the concentration 0.02 mg/ml and spectra was recorded on Shimadzu UV-visible recording spectrophotometer-UV-2100, supplied by Shimadzu co-operation, Japan).

### FT-IR Spectroscopy

To detect functional groups in the pure compound isolated from *S. apetala* leaf, FT-IR spectroscopy analysis was carried out in the Department of Chemistry, University of Mumbai (Shimadzu FTIR-4200 spectrometer, Shimadzu co-operation, Japan).

### Statistics

In this study, wherever applicable, all tests were performed in triplicates and data was expressed as mean ± standard deviation.

## Results

### Screening of anti-bacterial properties

Petroleum ether, acetone and water extracts were tested against Gram-positive as well as Gram-negative bacteria. Petroleum ether did not show activity against any bacteria. Water extract and acetone extract showed zones of inhibition against the Gram-positive as well as Gram-negative bacteria. The water and acetone extract has shown zones of inhibition ranging from 16-21mm. The water extract has shown anti-bacterial activity against four bacteria-*S. typhi, B. subtilis, Ps*.*auregenosa* and *E. Coli*, whereas acetone extract has shown zones of inhibition against all the bacteria except *Klebsiella aerogens* (Table-I).

**Table I.** Anti-bacterial potential of non-polar to polar solvent extracts in leaf of*S. apetala* plant.

| Bacterial strains | Anti-bacterial potential of extracts of *S. apetala* leaf (zones of inhibition in mm) |  |  |
| --- | --- | --- | --- |
|  | Petroleum ether | Acetone | Water |
| *Streptococcus pyogens* | -- | 20(±0.1) | -- |
| *Staphylococcus aureus* | -- | 21(±0.2) | -- |
| *Bacillus subtilis* | -- | 16(±0.1) | 18(±0.2) |
| *Staphylococcus epidermidis* | -- | 16(±0.3) | --- |
| *Escherchia coli* | -- | 18(±0.1) | 20(±0.1) |
| *Pseudomonas aeruginosa* | -- | 19(±0.2) | 20(±0.5) |
| *Salmonella typhi* | -- | 17(±0.3) | 18(±0.2) |
| *Klebsiella aerogens* | -- | -- | -- |

The MIC values for the acetone leaf extract (0.5, 1, 5 and 10 mg/ml) has shown inhibition against *S. typhi, B. Subtilis, Ps. auregenosa* and *E. Coli* (Table-II).

**Table II.** MIC of Acetone extract (Zones of Inhibition in mm)

| Zones of Inhibition (mm) against microorganisms | 10 mg/ml | 5 mg/ml | 1 mg/ml | 0.5 mg/ml |
| --- | --- | --- | --- | --- |
| *Streptococcus pyogens* | -- | -- | -- | -- |
| *Staphylococcus aureus* | -- | -- | -- | -- |
| *Salmonella typhi* | 16(±0.1) | 15(±0.2) | 12(±0.3) | 10(±0.3) |
| *Bacillus subtilis* | 15(±0.1) | 14(±0.1) | 12(±0.2) | 10(±0.3) |
| *Pseudomonas aeruginosa* | 15(±0.4) | 13(±0.1) | -- | -- |
| *Escherchia coli* | 15(±0.3) | 13(±0.3) | 11(±0.1) | 10(±0.1) |

To obtain appropriate solvent system for the separation and isolation of bioactive principles, column chromatography was done using either individual or mixture of solvents (non-polar and polar). In single solvent system, among all the solvents used (petroleum ether to methanol), diethyl ether and ethyl acetate have shown clear separation of bands (Table-III a). Diethyl ether and ethyl acetate have shown three bands whereas chloroform has shown four bands. The remaining solvents either showed trailing or no clear-cut separation of any band. The petroleum ether:ethyl acetate (7:3) resulted in five bands followed by acetone:chloroform(5:5) and petroleum ether:isopropanol(8:2) showing three bands. The solvent systems acetone:chloroform (8:2) and petroleum ether:methanol (7:3) separated only one band (Table III b).

**Table III a.** Column Chromatography usisng Single Solvent system.

| Solvents | No. of bands |
| --- | --- |
| Petroleum Ether | -- |
| Toulene | -- |
| Diethyl ether | 3 |
| Chloroform | 4 |
| Ethyl acetate | 3 |
| Isopropanol | -- |
| Acetone | -- |
| Methanol | -- |

**Table III b.** Column Chromatography using mixture of polar and non-polar solvents.

| Solvents | Solvent Proportion | No. of bands |
| --- | --- | --- |
| Petroleum ether: Ethyl acetate | 7 --- 3 | 5 |
| Acetone : Chloroform | 5 --- 5 | 3 |
| Acetone : Chloroform | 8 --- 2 | 1 |
| Petroleum ether : Isopropanol | 8 --- 2 | 3 |

Bioautography of all the chromatoplate showing clear cut separation of bands was carried out to observe active principles among separated bands. A single band obtained by the solvent system petroleum ether: methanol (7:3) was found to be active against *E. coli* and *B. subtilis* (Fig. II).

**Figure II.**
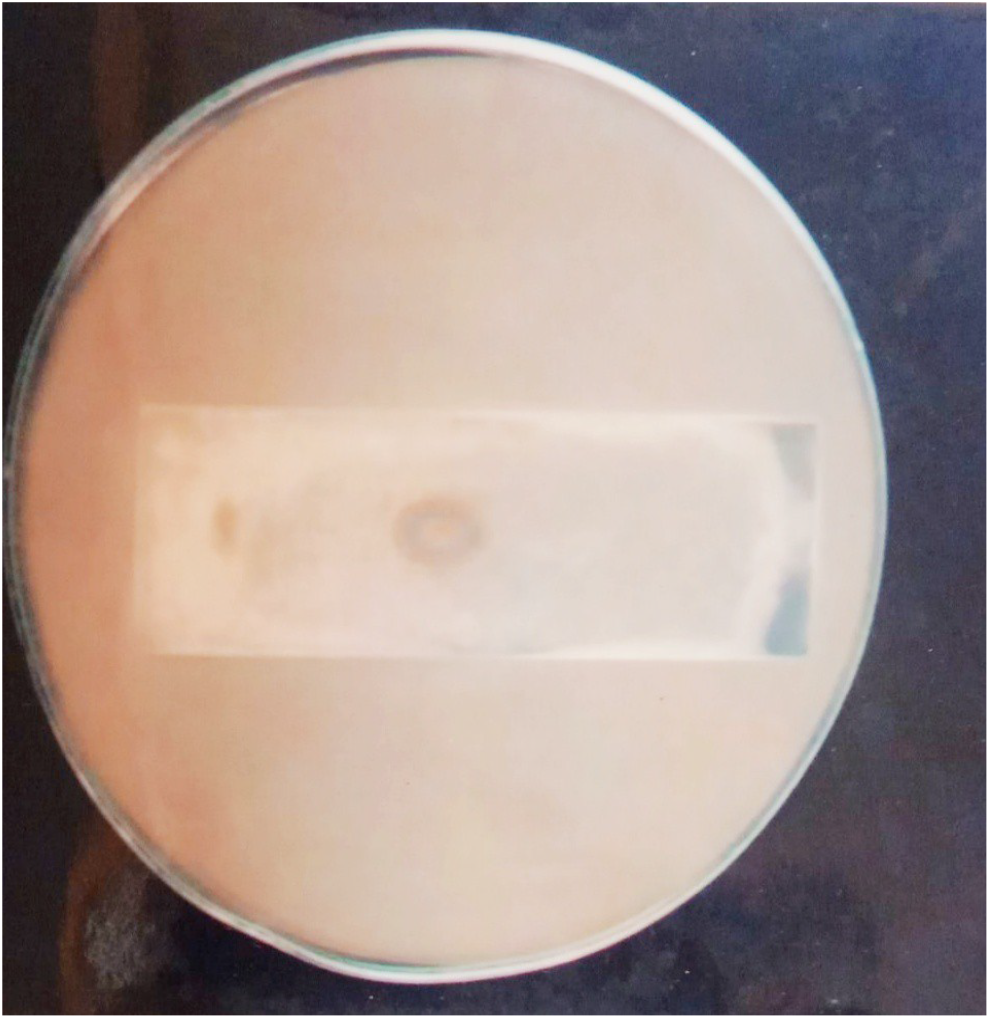
Petriplate showing Bioautography of the active fraction against *E*.*coli*.

None of the other band observed on chromatoplate developed by various solvent system showed anti-bacterial activity. Based on bioautography results, petroleum ether: methanol (7:3) solvent system was selected to isolate the bioactive compound by preparative column chromatography.

Purity of isolated compound was checked by TLC using 100% methanol as a solvent system where the prominent single spot was centrally developed on a TLC plate. Bioactivity of isolated compound along with standard antibiotics was studied against test microorganisms *E. Coli* and *B*.*subtilis* (Table-IV). The isolated compound showed the zones of inhibition 10 mm and 9 mm respectively against test bacteria which were better than ampicillin and Cefazolin. Similarly vancomycin, erythromycin and benzyl penicillium sodium showed comparable zones of inhibitions to that of isolated fraction which ranged between 12-14 mm.

**Table IV.** Activity of the given isolated compound with standard antibiotics.

| Standard antibiotics | *E.coli* | *B.subtilis* |
| --- | --- | --- |
| **Isolated compound** | **10** | **9** |
| Ampicillin | 8 | 8 |
| Cefazolin | 10 | 8 |
| Vancomycin | 13 | 13 |
| Erythromycin | 12 | 14 |
| Benzyl Penicilium sodium | 12 | 14 |

### Characterization and analysis of bioactive compound

The preliminary observations of active compound isolated by column chromatography like apperance, flame test and heating in a test tube were made and shown in Table-V. The compund was found to be solid, white or colorless, odorless crystals and completely soluble in water. A suity flame was observed in porcelain piece test while burning of the compound on copper wire exhibited green flame. Besides this, when strongly heated in a test tube instead of melting, compound was found to be charred.

**Table V.** Preliminary tests of the isolated compound.

| Serial number | Test | Observation |
| --- | --- | --- |
| 1 | Appearance:<br>i. Nature | Solid |
|  | ii. Colour | White or colourless |
|  | iii. Odour | No particular smell |
| 2 | Flame test |  |
|  | i. Porcelain piece test | Sooty flame |
|  | ii. Copper wire/Beilstein test | Burns with green flame |
| 3 | Heating in a test tube | Does not melt but chars on strong heating |
| 4 | Water solubility test | Water soluble |
| 5 | Detection of elements |  |
|  | i. Detection of N | No greenish blue colouration or precipitate |
|  | ii. Detection of Sulphur | No black precipitate |
|  | iii. Detection of halogen: |  |
|  | a. Test for halogen when both N and S absent | Thick white precipitate obtained (Halogen present) |
|  | b. Detection of individual halogen | Lower layer was colourless (Chloride present) |

#### Detection of elements

When an organic compound is fused with metallic sodium, the elements, ‘N’, ‘S’ and halogen are converted to corresponding sodium salts. On basis of this sodium fusion or Lassaigne’s test, element like ‘N’, ‘S’ and halogen were detected (Table-V). Nitrogen and Sulphur gave negative tests while halogen test was found to be positive with respect to chlorine.

#### Melting point

The uncorrected melting point of the compound was about 300 ^0^C.

#### UV-Vis Spectroscopy

The isolated compound showed a single absorption band at 208.5 nm (λmax) and the value for the absorbance of the compound was 0.740. This shows the compound comes in the UV spectrum. Consequently, it is colorless compound.

#### Infra Red Spectroscopy

FT-IR spectrum of the isolated compound was studied under two regions, the functional group region (4000-1300 cm^-1^ / 2.5-7.7 μm) and the fingerprint region (1300-900 cm^-1^ / 7.7-11.0 μm). The spectrum showed the presence of two absorption peaks at 1384 cm^-1^ and 1636 cm^-1^ regions. Besides this, one broad hump was observed at 3436 cm^-1^ region. The absorption in the region between 1950-1550 cm^-1^ characterised the ‘Carbonyl’ streching vibration. Absorption peaks located in the 1690-1600 cm-1 range arise from C=C and C=N stretching vibrations (Fig. III).

**Figure III.**
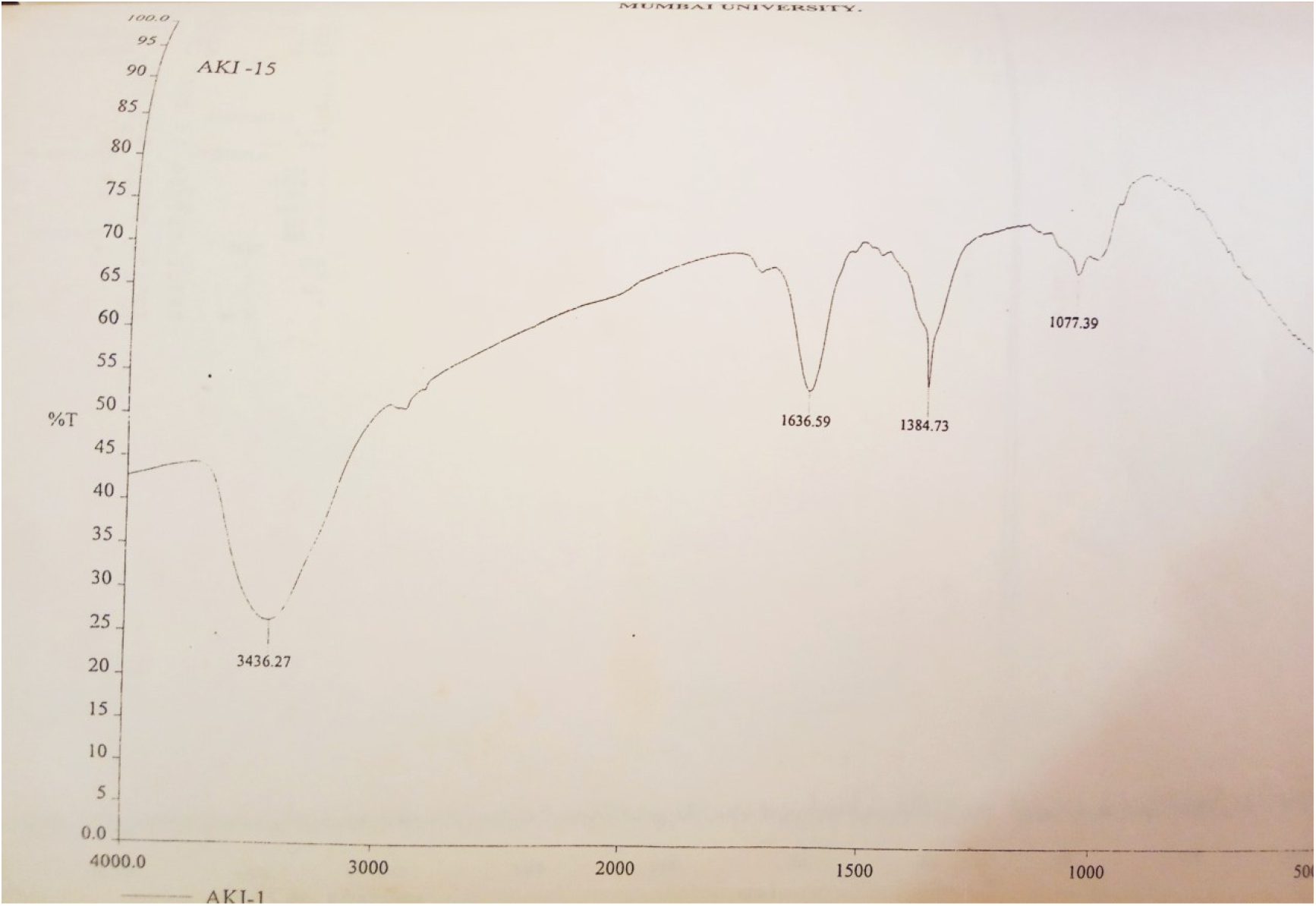
FT-IR spectrum of the isolated compound showing functional group region and fingerprint region.

## Discussion

The present study investigates the phytochemical profile of *S*.*apetala* leaf extracts obtained from the west coast of India and shows the potent anti-bacterial activity of *S. apetala* leaves. There exists some reports on anti-bacterial study of *S*.*apetala* leaves but the present study is different with respect to geographical location of plant collection, bacterial strains used and polarity of extraction solvents. In the current study, the bioassay-guided anti-bacterial fraction was isolated and characterized in contrast with earlier reports where primary screening for anti-bacterial activities were performed. This study along-with earlier reports validates the importance of *S*.*apetala* leaves as an important resource in traditional medicine for anti-bacterial remedies aimed at human pathogenic diseases (D. Jaimini et al., 2011; Patra et al., 2015).

This study has shown better results than the study carried out by Patra et al., where plant leaves were collected from the state of Odisha (Eastern coast of India). This also shows that the geographical location of the plant determines the anti-bacterial potency of the extract.

In order to isolate the bioactive components from the *S*.*apetala* leaves, solvent extraction was carried out by first selecting the appropriate solvent system using column chromatography with both single/mixture of solvents having varying polarity. Further, the purity and bioactivity of the isolated compounds was validated by TLC and against standard test antibiotics on different Gram-positive and Gram-negative bacteria.

In the present study, the leaf extracts were tested against different human pathogenic bacterial species like *S*.*pyogens, S*.*aureus, B*.*subitilis, E*.*coli, P*.*aeruginosa*, and *S*.*typhi* and potent anti-bacterial activity was observed against all these pathogenic species. MIC of crude extract(s) in the present study showed ability to inhibit the growth of pathogenic Gram-positive and Gram-negative bacteria at lower concentrations (0.5mg/ml) in both acetone and water-soluble extracts as compared to the Patra et al. study, where the MIC was found to be 2.5mg/ml and no activity established for pathogenic strains like *S*.*pyogens* and *S*.*typhi* (Patra et al, 2015). This result further validates the potent anti-bacterial actvity of *S*.*apetala* leaf extracts against a variety of pathogenic bacterial species and its potential as a traditional medicine for common bacterial diseases.

In order to gain more informtion on the appropriate solvent system and active compound(s)/fuctional group(s) present in the *S*.*apetela* crude extract, multiple solvent systems and chromatographic characterzations were carried out in the present study.

For the selection of the solvent system, preparative column chromatography was performed with either individual or a mixture of solvents (Polar to Non-Polar). For single solvent system, diethyl ether (3 bands), ethyl acetate (3 bands) and chloroform (4 bands) showed clear seperation and resolution of the bands.

In solvent combination system, petroleum ether:ethyl acetate (7:3) showed 5 bands while acetone: chloroform (5:5) and petroleum ether:isopropanol (8:2) showed 3 bands each while petroleum ether:methanol (7:3) showed one band respectively. This analysis thus indicated the efficacy and specificity of the various solvent system for isolation of the active compound(s) from crude extracts of *S*.*apetela* leaves.

Further TLC-bioautography method was carried out with the plates showing well separated bands. Here the petroleum ether:methanol (7:3) solvent system showed one band active against *E*.*coli* and *B*.*subtilis* validating the potential of *S*.*apetala* leaves as a potent anti-bacterial agent. The isolation of this anti-bacterial compound was carried out further by column chromatography and was confirmed as a single TLC spot.

Characterization of this organic compound was carried out using tests for solubility, melting point and elemental analysis via standard laboratory procedures.

UV-visible spectroscopy of this compound showed a single absorption band at 208.5 nm (λ max) indicating the compound to be UV-absorble in nature.

For obtaining more details about this compound, analysis via FT-IR spectroscopy was also performed (Pavia et al., 2008; Restiani and Nandiyanto, 2022). The FT-IR spectrum showed the presence of two absorption peaks at 1384 cm^-1^ and 1636 cm^-1^ regions. The absorption in the region between 1950-1550 cm^-1^ characterised the ‘Carbonyl’ streching vibration. Absorption peaks located in the 1690-1600 cm^-1^ range arise from C=C and C=N stretching vibrations.

The FT-IR analysis indicates the molecular absorption and transmission and it is possible that the potent anti-bacterial activty of *S*.*apetela* leaves may be due to the presence of aromatic and aliphatic groups (UV and FT-IR spectra) and detailed analysis with technique like ^1^H NMR spectroscopy will further elucidate the exact stucture and profile of this chemical compound.

Based on all the above investigations, it is clear that the acetone extract of *S*.*apetela* leaves show potent anti-bacterial activity against pathogenic Gram-positive and Gram-negative bacterial strains at low concentrations. TLC-bioautography of the chromatoplates also showed potent anti-bacterial activity against both *E*.*coli* and *B*.*subtilis* in the petroleum ether:methanol (7:3) solvent system. The use of such bioactivity-guided characterization of the active constituent(s) helps in their isolation and purification process. Further *in vivo* tests should be carried out to evaluate whether the potent anti-bacterial activity found *in vitro* is translated into *in vivo* activity.

The present study, therefore, elucidates the role of *S*.*apetela* leaves as a therapeutic agent in the treatment of bacterial infections and its use in traditional medicine.

